# Investigating the mechanism of chloroplast singlet oxygen signaling in the *Arabidopsis thaliana accelerated cell death 2* mutant

**DOI:** 10.1101/2024.01.25.577309

**Authors:** Matthew D. Lemke, Alexa N. Abate, Jesse D. Woodson

## Abstract

As sessile organisms, plants have evolved complex signaling mechanisms to sense stress and acclimate. This includes the use of reactive oxygen species (ROS) generated during dysfunctional photosynthesis to initiate signaling. One such ROS, singlet oxygen (^1^O_2_), can trigger retrograde signaling, chloroplast degradation, and programmed cell death. However, the signaling mechanisms are largely unknown. Several proteins (e.g., PUB4, OXI1, EX1) are proposed to play signaling roles across three *Arabidopsis thaliana* mutants that conditionally accumulate chloroplast ^1^O_2_ (*fluorescent in blue light* (*flu*), *chlorina 1* (*ch1*), and *plastid ferrochelatase 2* (*fc2*)). We previously demonstrated that these mutants reveal at least two chloroplast ^1^O_2_ signaling pathways (represented by *flu* and *fc2*/*ch1*). Here, we test if the ^1^O_2_-accumulating lesion mimic mutant, *accelerated cell death 2* (*acd2*), also utilizes these pathways. The *pub4-6* allele delayed lesion formation in *acd2* and restored photosynthetic efficiency and biomass. Conversely, an *oxi1* mutation had no measurable effect on these phenotypes. *acd2* mutants were not sensitive to excess light (EL) stress, yet *pub4-6* and *oxi1* both conferred EL tolerance within the *acd2* background, suggesting that EL-induced ^1^O_2_ signaling pathways are independent from spontaneous lesion formation. Thus, ^1^O_2_ signaling in *acd2* may represent a third (partially overlapping) pathway to control cellular degradation.

## Main text

To thrive while rooted to the ground, plants have acquired the ability to sense changes in their environment and acclimate. While plants employ diverse mechanisms to achieve such plastic physiologies, it is clear that they can use their chloroplasts (photosynthetic plastid organelles) to sense and respond to multiple types of abiotic and biotic stresses. This is due, in part, to chloroplasts being the site of photosynthesis. Such high energy metabolism is prone to producing reactive oxygen species (ROS), particularly under stress conditions such as drought, excess light (EL), salinity, and pathogen attack^1-3^. While these ROS can be toxic and lead to the damage of macromolecules in the chloroplast, they also act as signaling molecules^4^. One ROS in particular, singlet oxygen (^1^O_2_), is predominantly associated with chloroplasts (unlike superoxide and hydrogen peroxide, which are made in multiple cellular compartments) and produced primarily at photosystem II during photosynthesis^5. 1^O_2_ is known to trigger changes in nuclear gene expression (e.g., retrograde signaling), selective chloroplast degradation (i.e., chloroplast quality control), and cell death^6, 7^. Due to the extremely short half-life of ^1^O_2_ (< 1 μsec^8^), the bulk of this ROS is expected to remain within the chloroplasts in which it is made, necessitating the existence of signaling cascades. However, how ^1^O_2_ triggers such signaling and leads to these effects predominantly remains an open question in plant cell biology.

To study these signals, researchers have primarily relied on three *Arabidopsis thaliana* mutants that conditionally accumulate ^1^O_2_ in chloroplasts. The first characterized, *fluorescent in blue light* (*flu*)^9^, accumulates the tetrapyrrole chlorophyll precursor protochlorophyllide (Pchlide) in the dark. Upon a shift to light, this photosensitizing molecule leads to a rapid burst of ^1^O_2_ within chloroplasts that initiates a signal from the grana margins^10^. The *chlorina* (*ch1*) mutant cannot produce chlorophyll *b*, leaving the PSII reaction center without a protective antenna and sensitive to EL stress^11^. Under EL, *ch1* mutants produce a large amount of ^1^O_2_ at PSII within the grana core. Finally, a third mutant, *plastid ferrochelatase 2* (*fc2*), accumulates the chloroplast tetrapyrrole intermediate protoporphyrin IX (Proto) immediately after dawn^12^. Like Pchlide, free Proto also leads to the generation of ^1^O_2_ in the light, although the exact location of where the ^1^O_2_ is generated is unknown.

In all three mutants, the accumulation of chloroplast ^1^O_2_ leads to the induction of similar sets of nuclear marker genes, rapid bleaching of photosynthetic tissue, and eventual cell death^11-^ _13_. In the case of *fc2*, these signals also lead to selective^14^ and wholesale chloroplast degradation^12^, possibly depending on the bulk level of ^1^O_2_. Remarkably, these effects are not due to the toxicity of ^1^O_2_ per se, but to a genetically programmed response to its accumulation. In all three mutants, genetic suppressors have been identified through forward and reverse genetic approaches and many of these suppressor mutations can block signaling without reducing ^1^O_2_ levels_. 1_O_2_ signaling in *flu* can be blocked by mutations in *EXECUTOR1* (*EX1*) ^15^ and *EX2*^16^, which encode two chloroplast thylakoid-localized proteins. Signaling in *flu* is also blocked by mutations in *CRYTOCHROME1* (*CRY1*)^17^, which encodes a nuclear-/cytoplasmic-localized blue light photoreceptor. In *ch1*, mutations affecting *OXIDATIVE INDUCIBLE SIGNAL1* (*OXI1*) block ^1^O_2_ signaling^18^. OXI1 is a nuclear-localized Ser/Thr kinase, originally identified for its role in pathogen defense^19^. Finally, several mutations that block ^1^O_2_ signaling in *fc2* have been identified^12, 20, 21^. One mutation, *pub4-6*, affects the E3 ubiquitin ligase Plant U-Box 4 (PUB4), which may be involved (directly or indirectly) with the ubiquitination of proteins associated with the chloroplast envelope during photo-oxidative stress^22^. Such ubiquitination may be a mechanism by which photo-damaged chloroplasts are targeted for degradation^23^.

At first glance, these three mutants appear to generate the same ^1^O_2_ signaling pathway. They all induce programmed cell death (PCD) and regulate the expression of the same photo-oxidative stress nuclear marker genes. However, it is not known if these mutants represent one or multiple chloroplast signaling pathways. To this end, we recently used a meta-analysis of previously published whole transcriptome data sets of the three mutants to determine if they regulate the same set of genes^24^ due to generating the same ^1^O_2_ signal. We observed that they share a small core transcriptional response to ^1^O_2_ stress (36 genes), but maintain unique patterns. This result opened the possibility that these mutants use different ^1^O_2_ signals that report on specific stresses. Next, we combined the suppressor mutations described above with ^1^O_2_-producing backgrounds in which they were not originally isolated^24^. We then tested the ability of these suppressor mutations to block or alter ^1^O_2_ signaling in these new genetic backgrounds. Our results suggested that these mutants may represent two distinct ^1^O_2_ signaling pathways: one represented by *flu* (involving EX1/EX2 and CRY1), and one represented by *fc2* and *ch1* (involving PUB4 and OXI1). This was based on the observation that mutations that block signaling in *flu* (*ex1*/*ex2* and *cry1*) did not directly block signaling in *fc2*. The *cry1* mutation also did not block signaling in *oxi1*, while *ex1*/*ex2* were previously shown not to affect ^1^O_2_ signaling in this mutant^18^. In addition, genetic suppressors of *fc2* (*pub4-6*) and *ch1* (*oxi1*) did not affect ^1^O_2_ signaling in *flu*. However, *fc2* and *ch1* seemed to share the same pathway. The *pub4-6* mutation was able to block EL-induced PCD in *ch1*, and the *oxi1* mutation was able to block PCD in *fc2* adult plants (but not seedlings). Together, these results pointed to chloroplasts using at least two separate ^1^O_2_ pathways to induce PCD and retrograde signaling.

A fourth, less characterized mutant, *accelerated death 2* (*acd2*), has also been shown to accumulate ^1^O_2_, leading to spontaneous lesion formation in adult leaves^25, 26^. The ACD2 protein is needed to convert red chlorophyll catabolite (RCC) to primary fluorescent chlorophyll catabolite (pFCC) during chlorophyll catabolism^27^. In *acd2* mutants, the photosensitive tetrapyrrole RCC accumulates, which is likely responsible for the burst of ^1^O_228_. The site of ^1^O_2_ production in *acd2*, however, is not clear. ACD2 is expected to be active near PSII^29^, but also localizes to mitochondria under stress conditions^30^. The latter observation may explain why ^1^O_2_ has been shown to accumulate in *acd2* mitochondria^31^. Nonetheless, overexpression of *ACD2* has been shown to delay the hypersensitive response (HR) induced by *Pseudomonas syringae* infection, suggesting the regulation of tetrapyrrole catabolism may play a role in a pathogen defense response^30^. In citrus, an ACD2 homolog is a target of the *Candidatus* Liberibacter Asiaticus effector protein SDE15, and this interaction suppresses the plant’s hypersensitive response^32^. Consequently, ACD2-related ^1^O_2_ production may act as a mechanism by which plants can trigger PCD in response to pathogen attack.

Previously, it was shown that *acd2* mutants do not produce a *flu*-like signal, as *ex1*/*ex2* and *cry1* mutations were unable to block spontaneous cell death phenotypes in *acd2*^30^. To test if *fc2* may share a pathway with *acd2*, we previously generated an *acd2 pub4-6* double mutant and showed that it had delayed lesion formation^24^, suggesting that the ^1^O_2_ produced in *acd2* mutants triggers PCD through a PUB4-related mechanism. However, it is unknown if this is the same ^1^O_2_ pathway also shared with the *ch1* mutant.

To test this, here we generated an *acd2 oxi1* double mutant (using *acd2-2*^26^ and the *oxi1-1* T-DNA allele *GABI_355H08*^18^) and assessed lesion formation in adult leaves. As shown in **Fig. 1A and B**, lesions begin to spontaneously form in *acd2* mutants after 21 days in cycling light (16h light/8 hours dark) conditions with 125 μmol photons m^-2^ sec^-1^ white light at 21 °C. As before, the *pub4-6* mutation delayed this lesion formation at least up through 37 days. However, the *oxi1* mutation had no discernable effect on this phenotype, and *acd2 oxi1* mutants exhibited a similar number of lesions to *acd2*. Next, we tested the effect of the *acd2* mutation on photosynthesis by measuring maximum photosynthetic efficiency (F_v_/F_m_) across entire rosettes. As shown in **Figs. 1C and D**, *acd2* mutants begin to exhibit lower F_v_/F_m_ values after 24 days, indicating a degree of photosynthetic stress and loss of chloroplast function. This effect was completely reversed by the *pub4-6* mutation, but not by the *oxi1* mutation. Finally, we tested the final dry weight biomass of these mutants to assess what effect the spontaneous lesions in *acd2* have on the final yield. As expected, *acd2* mutants had significantly reduced biomass (dry weight) compared to wt (**Fig. 1E**). This was partly reversed by the *pub4-6* mutation, but not by the *oxi1* mutation. Notably, the *pub4-6* mutant did not lead to a general increase in biomass, as shown by the reduced biomass (compared to wt) of the single mutant. Together, this shows that the *pub4-6* mutation protects chloroplasts and maintains photosynthesis in *acd2* mutants to positively impact the final biomass yield of the pant. The *oxi1* mutation, on the other hand, had no measurable effect on any of these phenotypes. Thus, lesion formation in *acd2* may be distinct from EL-induced ^1^O_2_ signaling, which involves both PUB4 and OXI1 in wt and *ch1* mutants^11, 24^.

**Figure 1.**
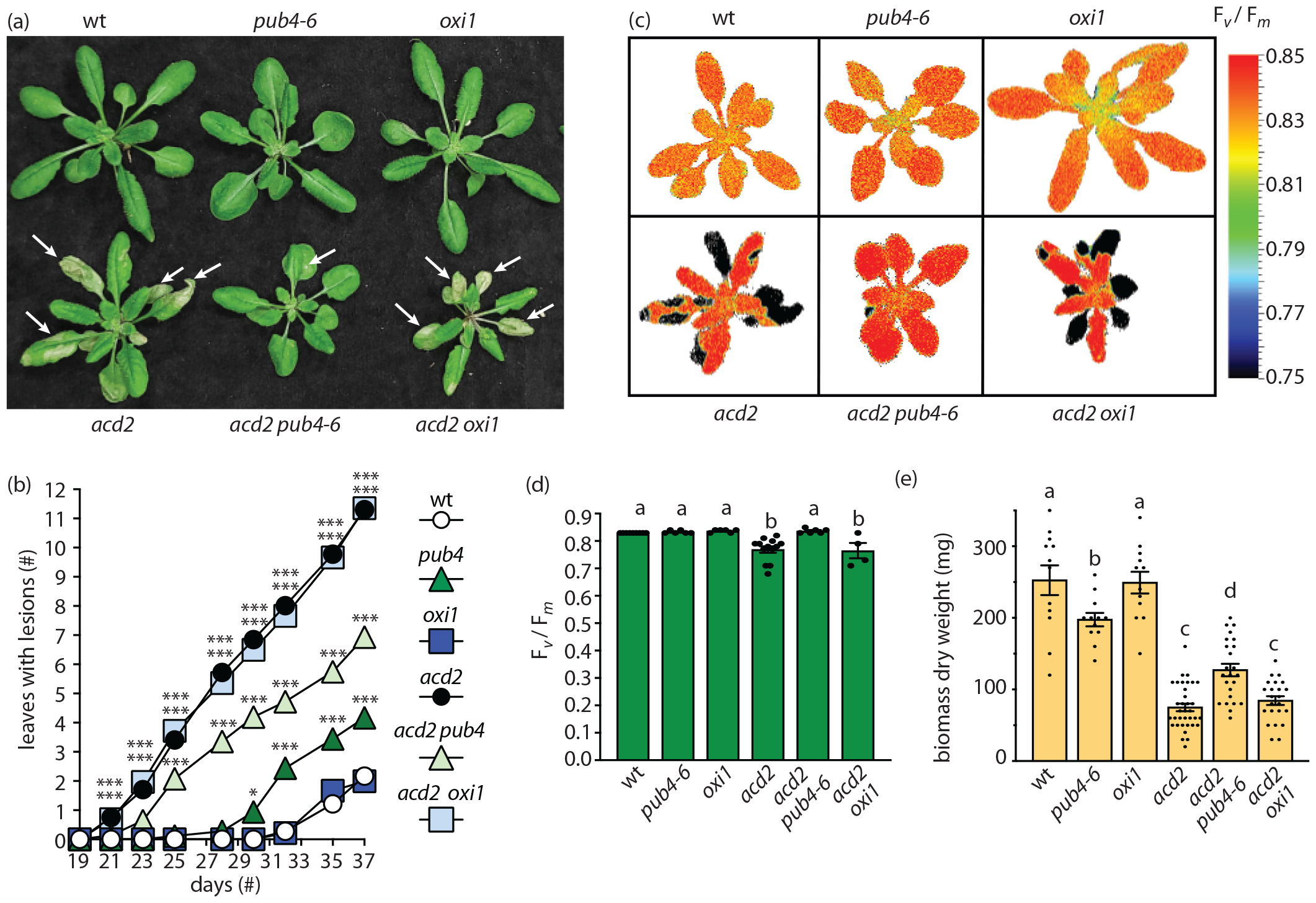
Assessing the effect of *pub4-6* and *oxi1* on lesion formation in *acd2*. **A)** Representative images of 24-day-old plants. White arrows indicate lesions. **B)** Assessment of mean lesion formation (number of rosette leaves with lesions) in plants between 19 and 37 days old (n ≥ 18 plants). **C)** Representative images of 24-day-old plants showing maximum photosynthetic efficiency (F_v_/F_m_) values. **D)** Mean F_v_/F_m_ values taken from plants in panel C (n ≥ 4 whole plant rosettes). **E)** Mean dry weight biomass (mg) from total aerial tissue of 57-day-old plants (n ≥ 12). Plant tissue was dried for three days at 65 °C before weighing. All plants were grown in cycling light conditions (16h light/ 8h dark) with 125 μmol photons m^-2^ sec^-1^ white light at 21 °C. Chlorophyll fluorescence measurements were conducted as previously described^37^. Statistical analyses were performed with a One-way ANOVA. In panel B, a Dunnett’s multiple comparisons post-test was used to test variation between genotypes relative to wt at each time point (* = *P* ≤ 0.05, ** = *P* ≤ 0.01, *** = *P* ≤ 0.001). In panels D and E, a Tukey’s multiple comparisons post-test was used to compare variation between genotypes. Different letters above bars indicate significant differences between genotypes (*P* ≤ 0.05). Error bars = +/-SEM. Closed circles indicate individual data points.

To test if this is the case in *acd2* mutants, we grew plants in cycling light conditions for 21 days and then exposed them to EL (1450-1550 μmol photons m^-2^ sec^-1^ white light) at 10 °C for 24h to specifically induce ^1^O_2_ stress^11, 18^. As shown in **Figs. 2A and B**, wt plants started to develop lesions after 24h. As previously shown^11, 24, 33^, the *pub4-6* and *oxi1* mutations delayed this lesion formation. Unexpectedly, the *acd2* mutants also had delayed lesion formation, with the *acd2 pub4-6* and *acd2 oxi1* double mutants having the least number of lesions. In the case of *pub4-6*, the effect was additive (*acd2* vs. *acd2 pub4-6, P*= 0.0304). Next, we measured the effect of EL on F_v_/F_m_ values. As shown in **Fig. 2C and D**, 6h and 24h of EL reduced F_v_/F_m_ in wt. As expected, *pub4-6* retained slightly higher values at 24h^24^. Surprisingly, *acd2* mutants had slightly increased F_v_/F_m_ values compared to wt, indicating an increased tolerance to EL. However, this was further increased in the *acd2 pub4* and *acd2 oxi1* double mutants and the effect was additive (*acd2* vs. *acd2 pub4*-6, *P*=0.0588; *acd2* vs. *acd2 oxi1, P*=0.0107). Together, these data suggest that *acd2* is not sensitive to EL stress, despite it potentially accumulating chlorophyll breakdown intermediates near PSII^29^. Furthermore, the EL stress signal is still disrupted in *acd2 pub4-6* and *acd2 oxi1*. This indicates that EL-induced lesions are unrelated to the spontaneous *acd2* lesions observed in permissive conditions, which may involve ^1^O_2_ signaling through a unique PUB4-related pathway.

**Figure 2.**
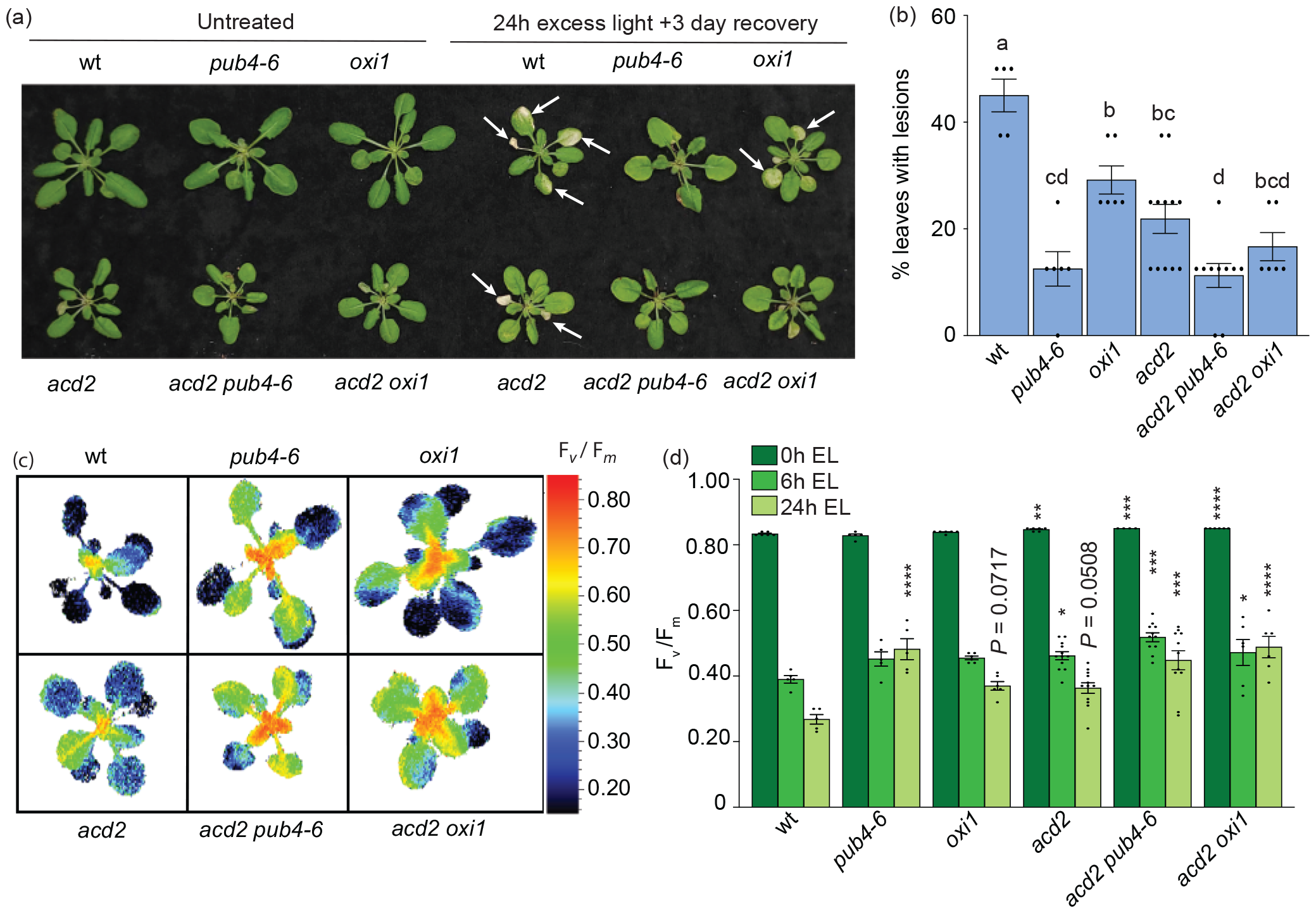
Assessing the tolerance of *acd2* mutants to excess light stress. Plants were grown for 21 days in cycling light (16h light/ 8h dark) conditions at 21 °C and then exposed to excess light (EL) at an intensity of 1450-1550 μmol photons m^-2^ sec^-1^ white light at 10 °C. **A)** Representative images of plants, either unexposed (left) or exposed to EL stress for 24 hours and allowed to recover for three days (right). White arrows indicate lesions. **B)** Mean % of leaves with lesions (ratio of leaves with observable cell death/healthy leaves) immediately after 24h EL exposure (n ≥ 5 plants). **C)** Representative images of stressed plants (immediately after 24h of EL) showing maximum photosynthetic efficiency (F_v_/F_m_) values. **D)** Mean F_v_/F_m_ values from whole plant rosettes after 0h, 6h, or 24h EL exposure (n ≥ 5 plants). F_v_/F_m_ values were obtained as previously described^33^. Statistical analyses were performed with a One-way ANOVA. In panel B, a Tukey’s multiple comparisons post-test was used to compare variation between genotypes. Different letters above bars indicate significant differences between genotypes (*P* ≤ 0.05). In panel D, a Dunnett’s multiple comparisons post-test was used to test variation between genotypes within a treatment relative to wt (* = *P* ≤ 0.05, ** = *P* ≤ 0.01, *** = *P* ≤ 0.001, **** = *P* ≤ 0.0001). Error bars = +/-SEM. Closed circles indicate individual data points.

Together, our results further define the ^1^O_2_ signaling pathways in plants and show that PUB4 and OXI1 represent two partially overlapping pathways in addition to the EX1/CRY1-dependent pathway identified in *flu* mutants (**Fig. 3**). While *oxi1* can block ^1^O_2_ signaling in *fc2* and *ch1, pub4-6* can block signaling in these mutants as well as the *acd2* mutant. As *acd2* may be producing ^1^O_2_ to combat pathogen or within mitochondria, it is tempting to conclude that PUB4 may be able to act more broadly in ROS signaling, possibly through immune responses that lead to PCD or hypersensitivity-like responses to pathogens. This is in line with recent reports linking PUB4 to basal defense pathways^34-36^ and tolerance to heat stress in the dark^33^. Such work underlines the complexity of chloroplast signaling, which may indicate how flexible these organelles are in sensing their environments and providing information for the cell. Further molecular characterization of PUB4 and these signaling pathways should help to reveal their signaling mechanisms.

**Figure 3.**
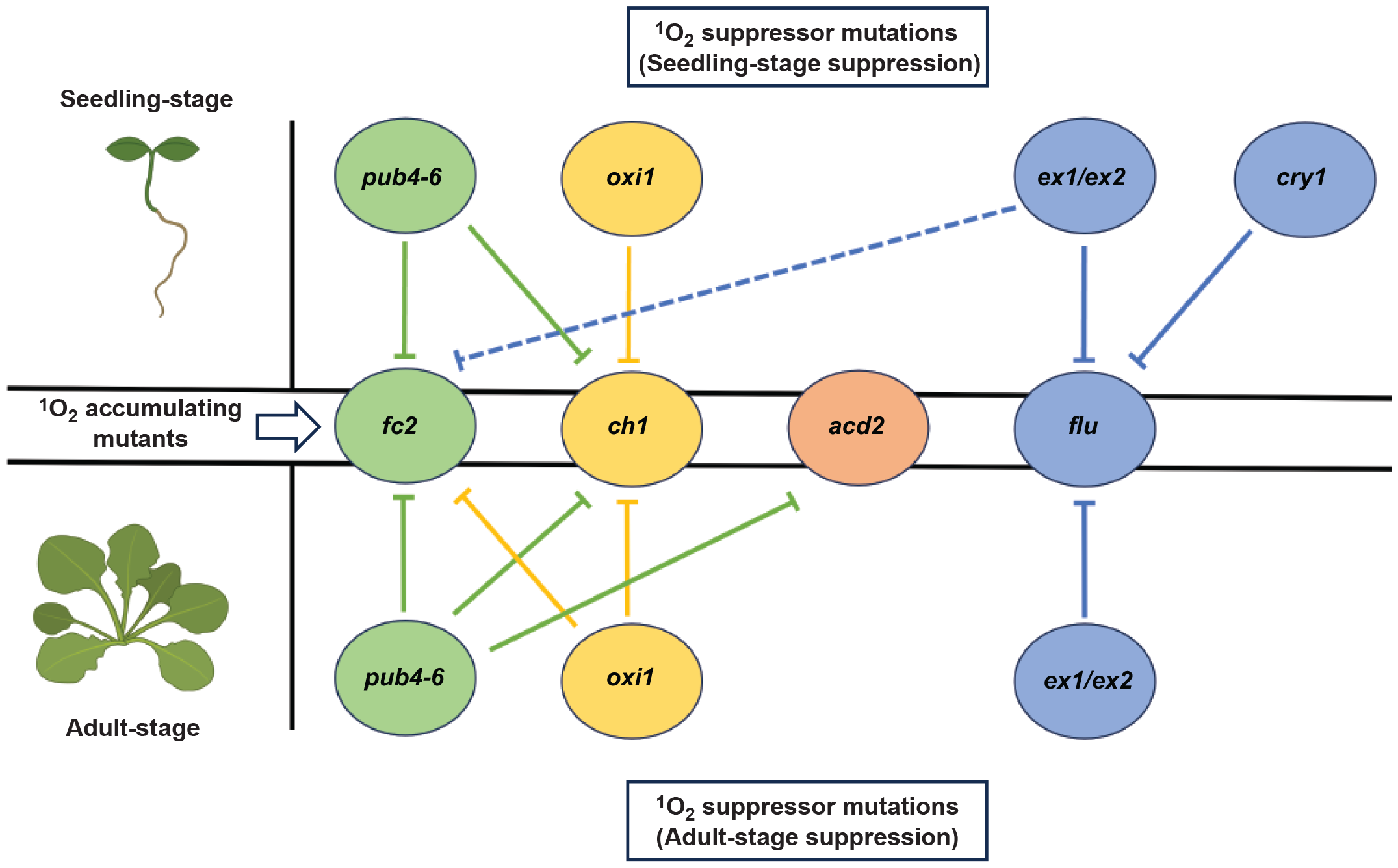
Differential suppression of singlet oxygen accumulating mutants by second-site mutations. A model summarizing the differential effects of secondary mutations on singlet oxygen (^1^O_2_) accumulating *Arabidopsis thaliana* mutants. Circles in the center row represent four ^1^O_2_ accumulating mutants: *plastid ferrochelatase 2* (*fc2*) (green), *chlorina* (*ch1*) (yellow), *accelerated cell death 2* (*acd2*) (orange), and *fluorescent in blue light* (*flu*) (blue). Circles on the top and bottom rows represent secondary mutations (*pub4-6, oxi1, ex1*/*ex2, cry1*) and their ability to block ^1^O_2_ signaling in the seedling (top) and adult stages (bottom). Circles sharing the same color as the ^1^O_2_ accumulating mutants indicate the mutant background in which these secondary mutations were first discovered. Solid lines and dashed lines indicate direct and indirect suppression of ^1^O_2_ signaling, respectively. This figure was created using online BioRender software (https://biorender.com/).

## Data availability statement

All data generated and analyzed during this study are included in this published article.

## Authors’ contributions

MDL and JDW planned and designed the research. MDL performed the assessment of photoinhibition and lesion formation in plants. ANA generated double mutants and confirmed by PCR. JDW conceived the original scope of the project and managed the project. MDL and JDW contributed to data analysis and interpretation, graphical visualization of the data, wrote the manuscript, and reviewed the manuscript. All authors approved the final version.

## Acknowledgments

The authors also wish to thank Sophia Daluisio (U of A) for technical assistance with genotyping.

## References

1. de Souza A, Wang JZ, Dehesh K. Retrograde Signals: Integrators of Interorganellar Communication and Orchestrators of Plant Development. Annu Rev Plant Biol 2017; 68:85–108.

2. Chan KX, Phua SY, Crisp P, McQuinn R, Pogson BJ. Learning the Languages of the Chloroplast: Retrograde Signaling and Beyond. Annu Rev Plant Biol 2016; 67:25–53.

3. Triantaphylides C, Havaux M. Singlet oxygen in plants: production, detoxification and signaling. Trends Plant Sci 2009; 14:219–28.

4. Mittler R, Zandalinas SI, Fichman Y, Van Breusegem F. Reactive oxygen species signalling in plant stress responses. Nat Rev Mol Cell Biol 2022; 23:663–79.

5. Triantaphylides C, Krischke M, Hoeberichts FA, Ksas B, Gresser G, Havaux M, et al. Singlet oxygen is the major reactive oxygen species involved in photooxidative damage to plants. Plant Physiol 2008; 148:960–8.

6. Woodson JD. Control of chloroplast degradation and cell death in response to stress. Trends in biochemical sciences 2022; 47:851–64.

7. Dogra V, Kim C. Singlet Oxygen Metabolism: From Genesis to Signaling. Front Plant Sci 2019; 10:1640.

8. Ogilby PR. Singlet oxygen: there is indeed something new under the sun. Chemical Society reviews 2010; 39:3181–209.

9. Meskauskiene R, Nater M, Goslings D, Kessler F, op den Camp R, Apel K. FLU: a negative regulator of chlorophyll biosynthesis in Arabidopsis thaliana. Proc Natl Acad Sci U S A 2001; 98:12826–31.

10. Wang L, Kim C, Xu X, Piskurewicz U, Dogra V, Singh S, et al. Singlet oxygen- and EXECUTER1-mediated signaling is initiated in grana margins and depends on the protease FtsH2. Proc Natl Acad Sci U S A 2016; 113:E3792–800.

11. Ramel F, Ksas B, Akkari E, Mialoundama AS, Monnet F, Krieger-Liszkay A, et al. Light-induced acclimation of the Arabidopsis chlorina1 mutant to singlet oxygen. Plant Cell 2013; 25:1445–62.

12. Woodson JD, Joens MS, Sinson AB, Gilkerson J, Salome PA, Weigel D, et al. Ubiquitin facilitates a quality-control pathway that removes damaged chloroplasts. Science 2015; 350:450–4.

13. op den Camp RG, Przybyla D, Ochsenbein C, Laloi C, Kim C, Danon A, et al. Rapid induction of distinct stress responses after the release of singlet oxygen in Arabidopsis. Plant Cell 2003; 15:2320–32.

14. Fisher KE, Krishnamoorthy P, Joens MS, Chory J, Fitzpatrick JAJ, Woodson JD. Singlet Oxygen Leads to Structural Changes to Chloroplasts During their Degradation in the Arabidopsis thaliana plastid ferrochelatase two Mutant. Plant and Cell Physiology 2022; 63:248–64.

15. Wagner D, Przybyla D, Op den Camp R, Kim C, Landgraf F, Lee KP, et al. The genetic basis of singlet oxygen-induced stress responses of Arabidopsis thaliana. Science 2004; 306:1183–5.

16. Lee KP, Kim C, Landgraf F, Apel K. EXECUTER1- and EXECUTER2-dependent transfer of stress-related signals from the plastid to the nucleus of Arabidopsis thaliana. Proc Natl Acad Sci U S A 2007; 104:10270–5.

17. Danon A, Coll NS, Apel K. Cryptochrome-1-dependent execution of programmed cell death induced by singlet oxygen in Arabidopsis thaliana (vol 103, pg 17036, 2006). Proc Natl Acad Sci U S A 2006; 103:18875-.

18. Shumbe L, Chevalier A, Legeret B, Taconnat L, Monnet F, Havaux M. Singlet Oxygen-Induced Cell Death in Arabidopsis under High-Light Stress Is Controlled by OXI1 Kinase. Plant Physiol 2016; 170:1757–71.

19. Rentel MC, Lecourieux D, Ouaked F, Usher SL, Petersen L, Okamoto H, et al. OXI1 kinase is necessary for oxidative burst-mediated signalling in Arabidopsis. Nature 2004; 427:858–61.

20. Alamdari K, Fisher KE, Sinson AB, Chory J, Woodson JD. Roles for the chloroplast-localized PPR Protein 30 and the “Mitochondrial” Transcription Termination Factor 9 in chloroplast quality control. Plant J 2020; 103:735–51.

21. Alamdari K, Fisher KE, Tano DW, Rai S, Palos KR, Nelson ADL, Woodson JD. Chloroplast quality control pathways are dependent on plastid DNA synthesis and nucleotides provided by cytidine triphosphate synthase two. New Phytologist 2021; 231:1431–48.

22. Jeran N, Rotasperti L, Frabetti G, Calabritto A, Pesaresi P, Tadini L. The PUB4 E3 Ubiquitin Ligase Is Responsible for the Variegated Phenotype Observed upon Alteration of Chloroplast Protein Homeostasis in Arabidopsis Cotyledons. Genes 2021; 12:1387.

23. Lemke MD, Woodson JD. Targeted for destruction: degradation of singlet oxygen-damaged chloroplasts. Plant Signal Behav 2022; 17:2084955.

24. Tano DW, Kozlowska MA, Easter RA, Woodson JD. Multiple pathways mediate chloroplast singlet oxygen stress signaling. Plant Molecular Biology 2023; 111:167–87.

25. Greenberg JT, Guo A, Klessig DF, Ausubel FM. Programmed cell death in plants: A pathogen-triggered response activated coordinately with multiple defense functions. Cell 1994; 77:551–63.

26. Mach JM, Castillo AR, Hoogstraten R, Greenberg JT. The Arabidopsis-accelerated cell death gene ACD2 encodes red chlorophyll catabolite reductase and suppresses the spread of disease symptoms. Proc Natl Acad Sci U S A 2001; 98:771–6.

27. Tanaka R, Kobayashi K, Masuda T. Tetrapyrrole Metabolism in Arabidopsis thaliana. Arabidopsis Book 2011; 9:e0145.

28. Pruzinská A, Anders I, Aubry S, Schenk N, Tapernoux-Lüthi E, Müller T, et al. In vivo participation of red chlorophyll catabolite reductase in chlorophyll breakdown. Plant Cell 2007; 19:369–87.

29. Sakuraba Y, Schelbert S, Park SY, Han SH, Lee BD, Andres CB, et al. STAY-GREEN and chlorophyll catabolic enzymes interact at light-harvesting complex II for chlorophyll detoxification during leaf senescence in Arabidopsis. Plant Cell 2012; 24:507–18.

30. Yao N, Greenberg JT. Arabidopsis ACCELERATED CELL DEATH2 modulates programmed cell death. Plant Cell 2006; 18:397–411.

31. Pattanayak GK, Venkataramani S, Hortensteiner S, Kunz L, Christ B, Moulin M, et al. Accelerated cell death 2 suppresses mitochondrial oxidative bursts and modulates cell death in Arabidopsis. Plant J 2012; 69:589–600.

32. Pang Z, Zhang L, Coaker G, Ma W, He SY, Wang N. Citrus CsACD2 Is a Target of Candidatus Liberibacter Asiaticus in Huanglongbing Disease. Plant Physiol 2020; 184:792–805.

33. Lemke MD, Woodson JD. A genetic screen for dominant chloroplast reactive oxygen species signaling mutants reveals life stage-specific singlet oxygen signaling networks. Front Plant Sci 2024; 14.

34. Desaki Y, Takahashi S, Sato K, Maeda K, Matsui S, Yoshimi I, et al. PUB4, a CERK1-Interacting Ubiquitin Ligase, Positively Regulates MAMP-Triggered Immunity in Arabidopsis. Plant and Cell Physiology 2019; 60:2573–83.

35. Wang Y, Wu Y, Zhong H, Chen S, Wong KB, Xia Y. Arabidopsis PUB2 and PUB4 connect signaling components of pattern-triggered immunity. New Phytol 2022; 233:2249–65.

36. Yu G, Derkacheva M, Rufian JS, Brillada C, Kowarschik K, Jiang S, et al. The Arabidopsis E3 ubiquitin ligase PUB4 regulates BIK1 and is targeted by a bacterial type-III effector. Embo J 2022; 41:e107257.

37. Lemke MD, Fisher EM, Kozlowska MA, Tano DW, Woodson JD. The core autophagy machinery is not required for chloroplast singlet oxygen-mediated cell death in the Arabidopsis plastid ferrochelatase two mutant. BMC Plant Biol 2021; 21:342.

